# Optogenetic Control of Activity in Descending *Tdc*^*2+*^ Neurons Modulates Motor Program Bias in the *Drosophila* Larval Locomotor System

**DOI:** 10.64898/2026.08.04.742787

**Authors:** William V. Smith, Karen Hibbard, Stefan R. Pulver

## Abstract

Motor systems must flexibly select between competing outputs while preserving stability of rhythmic outputs. In *Drosophila* larvae, the isolated central nervous system is capable of maintaining rhythmicity by generating multiple different fictive motor programs. The biogenic amines octopamine and tyramine are known to regulate larval locomotion, however, how the *tdc*^*2+*^ octopaminergic/tyraminergic system regulates motor program competition is not well understood. Here, we combine dual-colour calcium imaging and optogenetic manipulation to explore how *tdc*^*2+*^ neurons track, permit, and bias fictive motor activity in 3^rd^ instar *Drosophila* larvae. We find that *tdc*^*2+*^ activity in the larval ventral nerve cord is strongly coupled to motor neuron activity across multiple fictive behaviours, indicating that the system is recruited broadly across the motor repertoire. Optogenetic depolarisation of *tdc*^*2+*^ neurons increases motor root bursting and induces a robust fictive forward bias, whereas optogenetic hyperpolarisation suppresses or abolishes fictive rhythms and generates a short-lasting, post-inhibitory rebound in fictive activity. Spatially-restricted stimulation reveals that posterior abdominal activation is especially effective at promoting fictive forward activity. Separating VNC-residing from brain-residing *tdc*^*2+*^ populations further shows that activation of descending brain-residing *tdc*^*2+*^ projections is sufficient to recapitulate this forward bias. Finally, *tdc*^*2+*^ activation induces short-lived post-stimulation changes in motor programme probability, including transient elevation of competing fictive backward instantaneous frequency. Together, these findings suggest that *tdc*^*2+*^ neurons act as a permissive and biasing modulatory layer within larval motor circuits, linking adrenergic-like signalling to motor programme competition.

## Introduction

Motor systems must generate flexible but organised outputs from a finite set of circuit components. In *Drosophila* larvae, the isolated central nervous system spontaneously generates a diversity of fictive motor programmes functionally related to intact behaviour [1], [2]. These motor outputs are mutually exclusive, implying that larval motor circuits contain mechanisms that both generate diversity and regulate competition between alternative motor programmes [3]. Given their conserved role in altering excitability, rhythm stability, and activity thresholds, neuromodulatory systems are well positioned to shape motor competition [4], [5], [6], [7].

The *tdc*^*2+*^ system consists of a small set of well-characterised octopaminergic- and tyraminergic-releasing neurons; collectively the system provides a useful platform to explore the role of neuromodulatory systems in dynamically maintaining and shaping motor competition. The broad anatomy and locomotor-relevant properties of the larval *tdc*^*2+*^ system have been well characterised, but their role in actually shaping fictive dynamics has been unexplored. The larval system consists of distinct populations of octopaminergic and tyraminergic neurons across the central nervous systems, including brain-residing *tdc*^*2+*^ neurons that have local and descending projections into the ventral nerve cord (VNC) and VNC-residing neurons that send projections body-wall muscles [8], [9], [10], [11] (Figure 1). Previous work has demonstrated that *tdc*^*2+*^ neurons enable normal larval locomotion [12]. Further, genetic manipulations have demonstrated the role of octopaminergic and tyraminergic neuromodulation in coordinated locomotion [10], [13], [14] and behaviours associated with sensorimotor integration (e.g., learning) [15], [16], [17], [18], [19], [20], [21]. Despite a body of work, it remains unclear whether *tdc*^*2+*^ neurons merely modulate peripheral neuromuscular output via VNC-residing neuron’s projections or whether *tdc*^*2+*^ neuron activity is centrally coupled to fictive motor rhythms. Secondly, it is unclear how the *tdc*^*2+*^ system interacts with rhythmogenic networks controlling specific motor programs.

**Figure 1:**
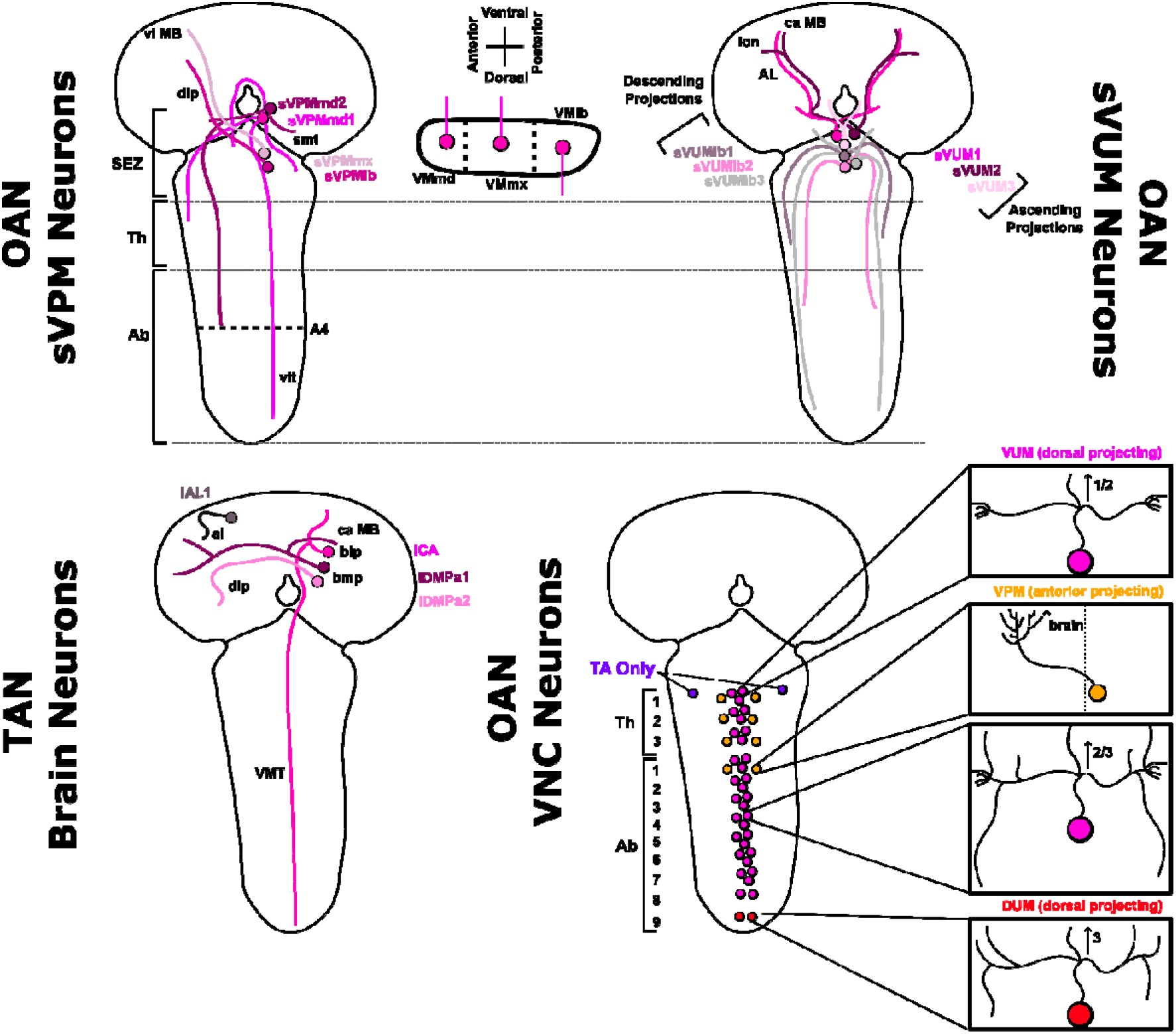
Octopaminergic and Tyraminergic Neurons in the *Drosophila* Larva Nervous System. General cell body position, defined name, and projection patterns for the octopaminergic (OAN) and tyraminergic (TAN) neurons as characterised in [8], [12]. The OAN ventral paired median (sVPM) neurons in the suboesophageal zone (SEZ) (only right-side shown) have contralateral brain projection and both ipsilateral and contralateral descending ventral nerve cord (VNC) projections. The projections of OAN sVPM neurons aligns to their anterior-to-posterior cell body position as shown beneath the sVPM map. The suboesophageal zone also harbours the suboesophageal ventral unpaired median (sVUM) octopaminergic interneurons which, unlike the VNC octopaminergic neurons, primarily function to dynamically modify sensory processing and context-dependent behaviours like arousal or odour discrimination [15]. The sVUM cluster can be classified into subsets with either ascending or descending projections. The vast majority of exclusively tyraminergic neurons are currently classified within the brain showing contralateral projections with interbrain connections, but some tyraminergic neurons do have VNC-wide descending projections. A set of ∼40 octopaminergic and 2 tyraminergic neurons reside exclusively within the VNC. As shown in the bottom right panel, OAN VNC neurons consist of ventral unpaired medial dorsal projecting, ventral paired medial anterior projecting to the brain, and dorsal unpaired medial dorsal projecting neurons. All of these neurons have local ascending projections 1-3 segments anteriorly of the segment which their somas reside. Importantly, only two neurons in [8], [12] were reported to exclusively manufacture tyramine and not octopamine; those neurons (shown in green) reside in the upper thoracic segment exclusively.

Here, we show that the 3^rd^ instar *Drosophila* larval *tdc*^*2+*^ system is tightly coupled to fictive motor output and can bias the network toward fictive forward locomotion. VNC *tdc*^*2+*^ activity tracks all fictive motor programmes, while *tdc*^*2+*^ neuron depolarisation increases motor bursting and induces strong fictive forward bias. In contrast, *tdc*^*2+*^ neuron hyperpolarisation suppresses or abolishes all fictive rhythms followed by a short-lasting post-inhibitory rebound in fictive activity. Forward bias can be uniquely driven by posterior stimulation of descending brain-residing *tdc*^*2+*^ neuron projections, rather than local VNC-residing *tdc*^*2+*^ neurons alone. Finally, brain-residing *tdc*^*2+*^ projection activation produces short-lived post-stimulation changes in motor program probability, including a transient elevation in fictive backward and long-lasting suppression in fictive forward instantaneous frequency. Together, these findings suggest that *tdc*^*2+*^ neuron act as a permissive and biasing modulatory layer over the larval motor network, linking adrenergic-like signalling to motor programme competition. Further, we instantiate how brain-residing *tdc*^*2+*^ neuron that project into the posterior VNC play a role in generation of fictive forward locomotion.

## Methods

### Animal Rearing & Dissection

*Drosophila* were raised at 18-25°C in incubators on an approximate 12-hour light-dark cycle. All experiments were conducted with feeding 3^rd^ instar *Drosophila* larva (2-3 days post hatching). The central nervous system (CNS) of *Drosophila* was isolated as reported in [1]. Activity of isolated CNSs was recorded with Baines External Saline (BES) submersion containing (in mM) 135 NaCl, 5 KCl, 2 CaCl_2_, 4 MgCl_2_, 5 TES, and 36 sucrose, pH 7.15 [22]. All CNS preparations were isolated from *Drosophila* larva using dorsal incision across the larva body using fine scissors with body walls pinned into a Sylgard-lined Petri dishes using fine tungsten wires (California Fine Wire, Grover Beach, CA). All projecting CNS nerves were cut, the CNS was removed, posterior nerves and connective tissues attached were pinned using fine tungsten wires (see [1]).

### Dual Colour Ca^2+^ Imaging of *Tdc*^*2+*^ and Motor Neurons

The calcium activity patterns within motor neurons and *tdc*^*2+*^ neurons [23] were assessed in isolated CNS feeding 3^rd^ instar *Drosophila* larva using a genetic construct that drove RCaMP1a expression in glutamatergic neurons (VGlut-LexA combined with 13XLexAOP-RCaMP1a balanced over CyO, Tb-RFP), while also containing UAS-GCaMP6f. This construct was then crossed to w[*]; P{w[+mC]=Tdc2-GAL4). Isolated CNS *Drosophila* were imaged using an OptoSplit Imaging system (Cairn Research Ltd, Kent, UK) with appropriate excitation and emission filters for RCAMP and GCAMP imaging. Preparations were imaged using an OLA light engine (Lumencor, OR, USA) and an ORCA-Fusion CMOS camera (Hamamatsu, Hamamatsu, Japan) with a XLUMPLFLN 20X dipping objective (Olympus, Tokyo, Japan). We imaged isolated dual-colour CNS *Drosophila* with BES using an X-light spinning disk confocal with a 70µm pinhole array (Crest Optics, Rome, Italy) for up to 30 minutes at 5fps with 150ms exposure. Images were collected using MicroManager (v2.0 gamma [24]) and post-processed using ImageJ [25], [26]. The channels displaying the calcium activity for motor neurons and *tdc*^*2+*^ neurons were overlayed using each segment’s nerve roots as a reference marker. All calcium signals were extracted using area-consistent regions of interest on nerve roots projecting from ventral nerve cord segments (T3-A6) (Figure 3Aii). Fluorescence traces were converted to ΔF/F by estimating a dynamic baseline for each ROI using a 20th-percentile filter over a 500-sample moving window. ΔF/F was calculated as (F − F_0_) / F_0_. Smoothed traces were generated from the ΔF/F signal using a second-order Savitzky–Golay filter with a 23-sample window. Fictive events were determined through using Hill’s Valley Analysis in DataView [3] and/or SciPy’s peak detection algorithm based on criteria reported below and in [1].

Visualisation of dual-colour calcium data was performed in DataView and Python3 using scripts available in paper-associated repositories. The overlaid average calcium activity trace in *tdc*^*2+*^ and motor neurons (Figure 3C) was performed using DataView’s event-scope view based on segregated fictive events detected by Hill’s Valley analysis across all preparations based on criteria reported in [1]. The average waveform of each event was calculated by DataView and extracted into Python for across preparation visualisation. All average waveforms were determined using a duration threshold of 8300ms. The peak time lag calculation (e.g. Figure 3D) was the difference in peaks from the average waveform of *tdc*^*2+*^ and motor neurons in each segment using SciPy’s peak detection algorithm (see repository code). Fictive events in calcium traces were determined using Hill’s Valley analysis and manually counted as reported in [1]. Peak and duration relation between *tdc*^*2+*^ and motor neuron calcium signals (e.g., Figure 3G) was performed using Hill’s Valley detection on each segment across all preparations with duration calculated as 20% on-off from the centre of the calcium signal. The correlation between *tdc*^*2+*^ and motor neuron calcium signals (as shown in Figure 3E) was conducted using Pearson Cross Correlation coefficient across all preparations through Python’s Scipy.Signal package (see repository code).

### *Tdc*^*2+*^ Optogenetics

The effect of endogenous *tdc*^*2+*^ activity on fictive motor activity was studied using extracellular electrophysiology on segmental nerve roots in isolated CNS from feeding 3^rd^ instar *Drosophila* larvae. Octopaminergic *tdc*^*2+*^ neurons were either depolarised using lines containing 20xUAS-CsChrimson-mVenus inserted at the attP18 landing site or hyperpolarized using lines containing UAS-GtACR1 inserted in VK00005 driven by w[*]; P{w[+mC]=Tdc2-GAL4. Across all conditions, the activity of thoracic nerve root (T3) and posterior abdominal nerve root (A6-A8) was measured by suction electrode recording (AM System Differential AC Amplifier Model (#59721, 1700), 1-100kHz cut-off frequency allowance). Borosilicate glass capillaries were pulled using an electrode puller (Narishege, Japan), and tapered tips were broken to produce suction electrodes that fit tightly around single nerve roots. Stimulation during the *tdc*^*2+*^ -CsChrimson conditions was performed with a downward-projecting red LED light source (Cairn Research Ltd., 620 nm, 0.58mW/cm^2^). The stimulation period was either consistent for 10, 20, or 40s or discontinuous with 0.01s pulses generated at intervals of 40Hz for ∼10s total pulsed stimulation. Stimulation during the *tdc*^*2+*^-GtACR1 conditions was performed with upward-projecting white light (∼2.5mW/cm^2^). The stimulation period was consistent for ∽15, 30, or 60s induced by manual instantiation. All whole-prep stimulation used full-prep illumination using a UPlanFLN Olympus 10x, 0.3NA lens.

Restricted illumination stimulation conditions involved restricting the downward-projecting red light source using a light-path restrictor with 0.6cm aperture and 0.16mW/cm^2^ power-to-area ratio. Restricted stimulation involved a LUMPlanF Olympus 40x, (0.8 NA) dipping lens, to prevent light scattering and thus unintended illumination, focused on posterior abdominal, anterior abdominal, thoracic, and suboesophageal ganglion regions (see Figure 6). All conditions were matched with control stimulation where illumination of the left-flank of the preparation (< 1cm) was conducted during nerve root recording.

Optogenetic manipulation of brain-residing and VNC-residing *tdc*^*2+*^ neuron was conducted via restricted GAL4 expression. *CsChrimson* was solely expressed in VNC-residing *tdc*^*2+*^ neurons by crossing flies containing 20XUAS>dsFRT>CsChrimson, tsh-LexA, 8xLexAopFlp attP40/CyO, Tb-RFP with w[*]; P{w[+mC]=Tdc2-GAL4. In contrast, *CsChrimson* was solely expressed in brain-residing *tdc*^*2+*^ neuron by crossing 20XUAS-CsChrimson-venus in attP18;tsh-LexA, pJFRC20-8XLexAop2-IVS-GAL80-WPRE attP40/ CyO, Tb-RFP; MKRS/TM6B flies with w[*]; P{w[+mC]=Tdc2-GAL4 animals. Expression patterns were verified using confocal microscopy shown in Supplementary Figure 2, 3.

When instantaneous frequency changes as a result of optogenetic stimulation were assessed, 10 second averaged windows of each fictive behaviour’s frequency were reported. Instantaneous frequency was calculated through peak detection within DataView [27]. Specifically, within instantaneous frequency average plots (e.g., Figure 8D), the frequency and time were normalised. Normalisation was scaled within each preparation for each specific behaviour where 1 represents the maximum instantaneous frequency found for that particular fictive behaviour within that particular preparation stimulation period. Instantaneous fictive-frequency traces were organised into chronological repeats defined by each optogenetic stimulation epoch, with adjacent pre-stimulation and post-stimulation epochs assigned to the same repeat; each segment epoch was then independently time-normalised to an equal-width interval — 0–1, 1–2 and 2–3, respectively — interpolated onto a common grid, normalised within each repeat behaviour, smoothed with a Gaussian filter, and plotted as individual repeat traces with the across-repeat mean overlaid.

All electrophysiology recordings and light stimulations were made using LabChart 8.0 software (AD Instruments, Colorado Springs, CO) and post-processed in Spike2 (Cambridge Electronic Design, Cambridge, UK) and Python3. Post-processing involved rectification and smoothing using a consistent 0.2s time interval. All rectified and smoothed traces were imported into DataView where event peaks and 20% on-off duration were determined by Hill’s Valley analysis. Visualisation and further analysis were conducted in Python3 (see repository code).

### Classifying Fictive Activity

Fictive activity was classified based on detecting peak in motor neuron calcium signals and using strict definitions of intersegment peak activity. The motor dynamics within the isolated VNC preparation were classified using Hill’s Valley thresholding to determine peaks within motor activity in each abdominal and thoracic ganglion segment alongside established definitions for each fictive behaviour [1], [2], [3] (Figure 2). Succinctly, fictive forward waves and fictive backward waves were determined by the existence of sequential Hill’s Valley-determined peaks proceeding from either posterior-to-anterior or anterior-to-posterior in all segments, respectively. Fictive headsweeps were determined by computing a difference trace (left – right) in the T3 segment and determining anterior asymmetric activities as peaks or troughs that exceeded ±10% absolute threshold difference. Finally, anterior symmetric (i.e., anterior burst) or posterior symmetric (i.e., posterior burst) were determined by peak detection of the left and right trace in T3-A2 and A6-8, respectively. Some anterior burst and posterior burst fictive activities are in varying numbers of anterior and posterior segments thus any peak detection in both the left and right trace within 5s of each other was determined as symmetric activities of each definition. In the calcium dataset, Hill’s Valley peak detection was performed after baseline smoothing within DataView [3]. In the electrophysiology dataset, raw electrophysiological traces were smoothed in Spike2 thereafter DataView was used to determine peaks in the anterior and posterior trace akin to the calcium dataset analysis pipeline. Specifically, the determination of fictive forward vs backward activity was determined by the minimum peak-to-peak time between the initiating-vs-terminating segment. Specifically, if the peak time difference between T3-to-A8 was shorter than the peak time between A8-to-the-next-T3 peak then the activity peaks were classified as fictive backward events rather than fictive forward events. If the activity peak differences exceeded 3s, the activity peaks were either reported as anterior bursting or posterior bursting events in T3 or A8, respectively. In nerve root recordings, posterior bursts tend to be sharper for fictive backward rather fictive forward activity patterns, an auxiliary definition used in instances of uncertainty. In cases of significant ambiguity, the fictive events were not recorded. Manual corrections – deletions or additions of peaks – were performed, when necessary, in line with the strict fictive definitions highlighted above.

**Figure 2:**
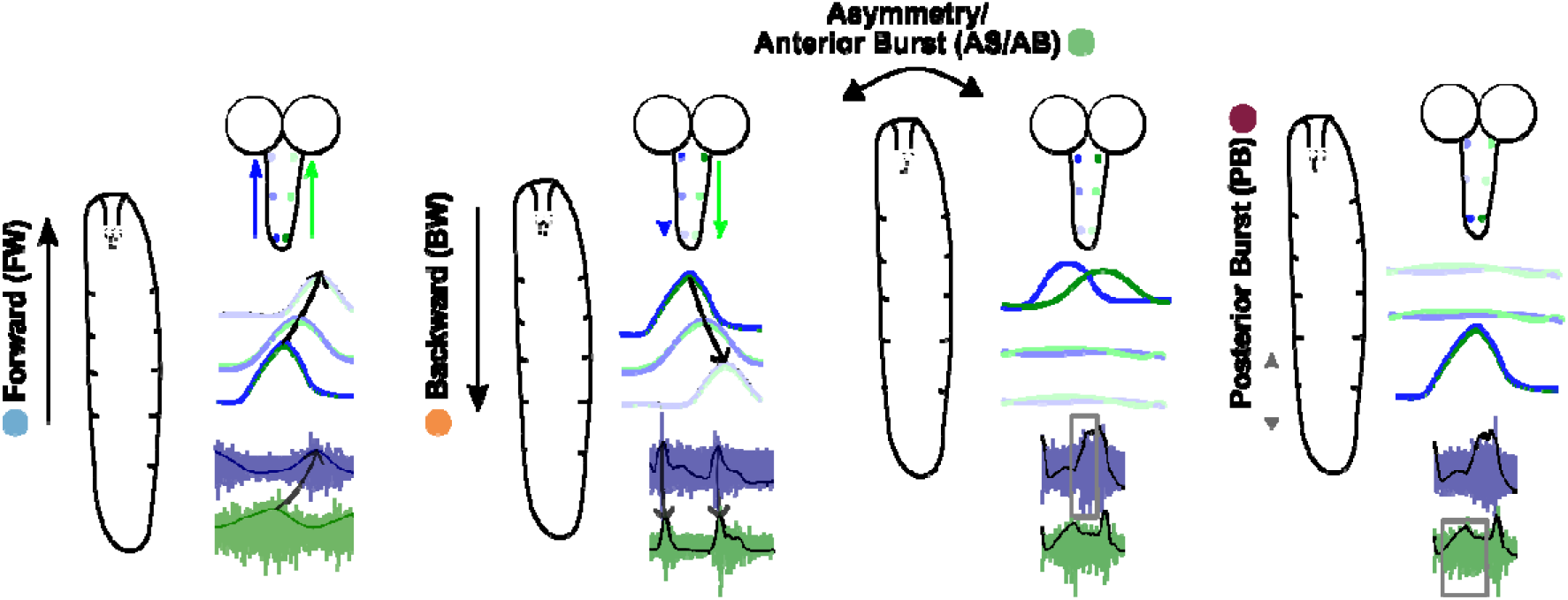
Fictive Behavioural Categories. The intact behaviour of the *Drosophila* larva is classified by different motor neuron activity sequences within the isolated ventral nerve cord (as originally noted in [1], [2]). Fictive activity patterns are shown alongside analogous intact behavioural patterns. Forward crawling consists of segment contractions in a posterior-to-anterior direction induced by a posterior-to-anterior sequence of motor neuron activity and temporal sequence of motor root bursting. Backwards crawling consists of segment contractions in an anterior-to-posterior direction induced by an anterior-to-posterior sequence of motor neuron activity and temporal sequence of motor root bursting. Headsweep behaviour – angular sweeping of the anterior regions of the intact animal – is induced by asymmetric activation of either the left or right side of motor neurons in the thoracic ganglion and exclusive bursting in one lateral anterior nerve root (see grey box). Head crunches/raises are likely induced by anterior symmetric activation of motor neurons in the thoracic ganglion and temporal sequence of motor root bursting exclusive bursting the anterior nerve roots. Posterior crunches consist of posterior segment contractions induced by symmetric activity in the motor neurons in the posterior segments of the abdominal ganglion and exclusive bursting in posterior nerve roots (see grey box). All subsequent calcium imaging and electrophysiology analysis categorises sequential motor activity in either of the six fictive patterns: fictive forward (FW), fictive backward (BW),anterior asymmetric (AS), turns (TR), anterior bursting (AB), or posterior bursting (PB) events

The duration of total fictive activity was determined through the same Hill’s Valley peak detection algorithm used to initially categorise fictive activity patterns. Specifically, the total duration of fictive activity within a condition time was calculated as the 80% margins before and after peaks in calcium activity from all segments that exceeded 10% standard deviation above the baseline signal. Silence was determined by the total time of each condition’s recording period subtracted by the duration of total fictive activity. Fictive overlap was defined as the existence of the start, or progression, of a fictive behavioural program in the network before the definitional completion of a prior fictive program. For instance, a fictive forward wave would be overlapping a fictive backward wave if the A8 initiation peak of the fictive forward wave occurring before the T3 completion peak of the fictive backward wave. Exclusively fictive wave motor programs were considered in the definition of overlapping fictive events.

### Analysis, Visualisation, & Statistical Evaluation

All calcium signal and electrophysiology nerve root data were initially analysed in DataView (version 12.2.2)[28] using Hill’s Valley Analysis for peak detection and Event-Triggered Scope View for averaging waveforms. All data were visualised in Python3 using custom scripts available in the repository supplied by the authors. All statistical tests were performed using packages within Python.

### Sustainability Statement

Using the carbon calculator WillCO_2_st [29], [30], we estimate that experimental work presented here generated 645.19g CO_2_e from the South Scotland UK National Grid.

## Results

### *Tdc*^*2+*^ VNC Neuron Activity is Correlated with Motor Neuron Activity across all Fictive Motor Programs

The isolated *Drosophila* central nervous system (CNS) spontaneously generates motor rhythms (denoted as “fictive” to distinguish from intact motor programs) in the thoracic and abdominal segments that can be measured via calcium indicators expressed in neuronal populations [1]. Previous work has characterised the features and relationship between fictive activity patterns using different genetically-encoded calcium indicators [1], [2], [3], [31]. Succinctly, isolated CNS preparations display fictive forwards and backwards motor programmes characterised by anterior-to-posterior or posterior-to-anterior progression of activity in motor neuron segments, respectively. In addition, preparations also can display fictive bilaterally symmetric activity in the anterior thoracic and posterior abdominal segments indicative of head raising or tail grounding behaviours, respectively. Finally, preparations can display bilaterally asymmetric activity in the anterior thoracic indicative of headsweep-like behaviours in intact *Drosophila* larva (Figure 3Ai). Octopamine (OA) is synthesised and released from neuronal subpopulations in the suboesophageal ganglion, ventral nerve cord (VNC), and brain expressing *tdc*^*2+*^ [12], [32]. Given both chronic and acute manipulations can have a profound effect on locomotion, we first evaluated if activity of OA neurons in the VNC was correlated with fictive motor rhythms. There exist subpopulations of distinct octopaminergic neurons that line the ventromedial (T3-A6) axis of the CNS [12]. Through expressing different calcium indicators in VNC motor neurons (VGlut, RCaMP1a) and ventromedial *tdc*^*2+*^ neurons (TDC^2+^, GCaMP6f) (Figure 3Aii), we explored the activity relationship between VNC *tdc*^*2+*^ neurons and motor neurons during spontaneous fictive activity bouts. The calcium activity of motor neurons and *tdc*^*2+*^ neurons in each VNC segment showed high correlation across different fictive motor programs (Figure 3B).

**Figure 3:**
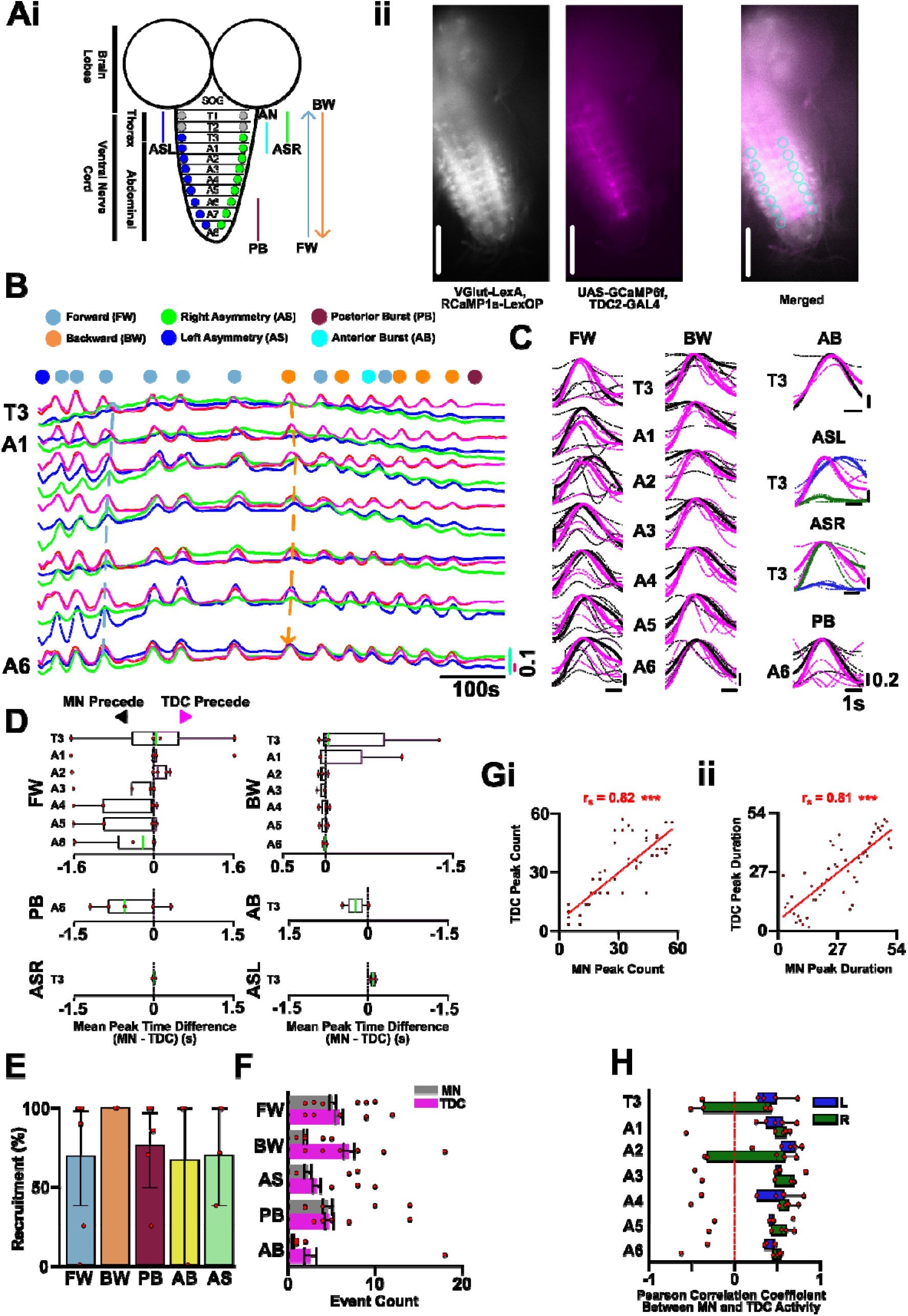
High Activity Correlation between *tdc*^*2+*^ VNC and Motor Neurons in the Same Segment and Fictive Pattern. (**Ai**) The isolated ventral nerve cord (VNC) demonstrates sequences of motor neuron activity resembling activity patterns underlying locomotion in intact animals (i.e., “FW” = fictive forward, “BW” = fictive backward, “AB” = anterior burst, “PB” = posterior burst, “ASR” = asymmetric right, “ASL” = asymmetry left). (**Aii**) Intrinsic calcium activity of motor neurons (MN) and octopaminergic neurons (*tdc*^*2+*^) is evaluated through dual calcium indicator expression at nerve root regions of interest (blue circles) T3 -A6. Dots above the traces indicate the initiation of the different fictive behaviours (light blue, fictive forward; orange, fictive backward; yellow, anterior symmetric burst; pink, posterior burst; blue, left anterior asymmetry; green, right anterior asymmetry). Lines indicate baseline-correct calcium fluorescence values (green/blue = right/left MN in each segment, magenta/red = right/left *tdc*^*2+*^ VUM in each segment) (**B**) Exemplar calcium trace of MN (left = blue, green = right) and *tdc*^*2+*^neurons (left = magenta, right = red) demonstrating strong association between MNs and Tdc2+ neuron activity within each ventral nerve cord (VNC) segment. Blue and orange arrow indicates the progression of an exemplar fictive forward and backward wave. (**C**) Averaged calcium image traces of *VGlut RCaMP1a* (black), *Tdc2 GCaMP6f* (magenta) for fictive forward (FW, black for left and right MN average), backward (BW), anterior burst (AB), left asymmetry (blue, ASL), right asymmetry (green, ASR), and posterior burst (PB) activity. Bold traces represent average across all preparations (N = 7) with each individual prep average calcium trace in non-bold, dashed lines. The calcium signal amplitudes are normalised within each fictive behavioural class. The y-axis for each segment are the time frame relative between the TDC and MN trace of that segment, not standardised across segments thus fictive wave progression is not shown. (**D**) The time difference between the peaks in the MN and TDC calcium trace within each preparation, across fictive patterns and ventral nerve cord segment (T3 - A6). Negative average peak time differences occur when the average prep MN trace proceeds the TDC signal in the segment, positive average peak time represents TDC proceeding MN signals in the segment. Green line represents the average peak time difference across all preparations in the most distal VNC segments. (**E**) The % event count where a motor neuron fictive pattern exists alongside a corresponding TDC activity pattern across different fictive activity categories (with 95% confidence interval bars). Recruitment % represents the number of MN and TDC events divided by the total fictive events. (**F**) The count of fictive activity patterns demonstrated by MN and TDC VNC populations. Note in many fictive activity patterns there is an excess of TDC exhibitions compared to motor fictive patterns. (**G**) MN and TDC signals pooled across segments and preparations showing the Spearman’s Rank for (**i**) peak count comparison and (**ii**) 20% on-off peak duration comparison. (**H**) Pearson correlation coefficient for the MN and TDC traces of each segment within all preparations for the left (blue) and right (green) axis of the VNC. Comparisons followed by post-hoc statistics were conducted with p-values: < .5 (*), < .01 (**), <.001 (***) shown.

By parsing all instances of fictive events across dual-colour preparations (N=7), we can see *tdc*^*2+*^ and motor neuron are regularly active concurrently within each segment (Figure 3C). Specifically, if we take the average calcium activity trace for all segments for motor neurons and *tdc*^*2+*^ neurons across fictive activity patterns, we see motor neuron *tdc*^*2+*^ neurons activity has a consistent similarity and phase locking. The strong activity association between motor neurons and *tdc*^*2+*^ neurons is further apparent when comparing activity peaks between each neuronal population in each segment per each unique fictive behaviour (Figure 3D). The temporal relationship between motor neuronal and *tdc*^*2+*^ neurons showed variations depending on the fictive motor program. At the initiating segment for fictive activity, motor neuron activity on average preceded *tdc*^*2+*^ neuron activity for fictive forward waves (462±704 ms at A6, N = 41 instances), posterior burst (449±614 ms at A6, N = 31 instances), and anterior burst (235±352 ms at T3, N = 2 instances) events. In contrast, *tdc*^*2+*^ neuron activity preceded motor neuron activity in initiating segments for fictive backward (432±786 ms at T3, N = 7 instances), right asymmetric (9±38 ms at T3, N = 5 instances), and left asymmetric (99±57 ms at T3, N = 2 instances) events. Importantly, the time phase relationship between motor neurons and *tdc*^*2+*^ neuron activity at initiation is not always held consistently in all ganglion segments as the activity pattern progresses through the VNC. For example, as fictive forward motor activity progresses to the anterior regions (A2 - T3) of the CNS, average *tdc*^*2+*^ neuron peak activity now precedes motor neuron activity (Figure 3C, D). In short, while there is some inter-prep variability between the temporal organisation of motor neuron and *tdc*^*2+*^ neuron peak activity, there is a consistent co-current activity between the activity of these two neuron subpopulations in all fictive behaviours; in all fictive patterns, there is a system-wide mirror-image of activity across all segments even if the lag relationships show intersegmental differences.

*Tdc*^*2+*^ neurons and motor neurons exhibit fictive-pattern-specific-activity correlation. The majority of fictive motor events across preparations showed commensurate *tdc*^*2+*^ neuron activity events. Across all preparations, 69±47% of fictive forward waves, 70±77% of anterior asymmetric events, 76±39% of posterior burst events, and 100±0.0% fictive backward waves showed commensurate *tdc*^*2+*^ activity in a close time window (< 5s) and same segment-to-segment relationship that defines the fictive motor programs (Figure 3E). There were instances of fictive-like *tdc*^*2+*^ activity patterns that occur independent on any co-activation with fictive motor patterns. Across all preparations there were 7.8% more backwards-like activity patterns in *tdc*^*2+*^ neurons than backward fictive motor activity patterns despite 100% recruitment per preparation (Figure 3F). Nonetheless, per preparation, there were 8.7%, 5.4%, 1.8%, 2.1% excess *tdc*^*2+*^ recruitment for anterior burst, anterior asymmetric, fictive forward, and posterior burst activity, respectively, the event counts between motor neuron and *tdc*^*2+*^ neurons activity were not significantly different.

Independent of fictive events, the activity in motor neurons and *tdc*^*2+*^ neuron pools per segment are correlated. Specifically, the number of motor neuron and *tdc*^*2+*^ neuron calcium activity peaks (r_s_ = 0.82, p < .001) (Figure 3Gi) and their duration (r_s_ = 0.81, p < .001) (Figure 3Gii) of activity within each segment were correlated. Pearson correlation between motor neuron and *tdc*^*2+*^ neuron calcium activity per segment demonstrated positive correlation (r = 0.4; Figure 3H) but with considerable intersegmental and lateral variability (Fisher’s Z Score = 0.44 with 0.27 variance). Overall, activity of motor neurons and *tdc*^*2+*^ neurons are highly correlated.

### *Tdc*^*2+*^ Neurons show Command-like Properties for Fictive Locomotor Activity

We next evaluated if manipulating intrinsic *tdc*^*2+*^ system activity can alter motor competition. We expressed either a depolarising red-light-gated-channel (*CsChrimson*) or a hyperpolarising green-light-gated channel (*GtACR1*) in *tdc*^*2+*^ neurons to optogenetically manipulate the intrinsic adrenergic-like system and evaluate motor neuron activity using nerve-root electrophysiology. *Drosophila* larvae reared on retinol-based food facilitates the appropriate folding of the opsins and thus constitutes the experimental line whereas *Drosophila* reared on food devoid of retinol, and thus appropriate opsin folding and insertion, act as the experimental control [33], [34].

Firstly, we optogenetically depolarised *tdc*^*2+*^ neurons expressing *CsChrimson* by continuous red-light illumination (10, 20, 40s) and recorded motor root activity from left and right T3 and A6 VNC ganglia (Figure 4A). Optogenetic depolarisation of *tdc*^*2+*^ neurons induced increased A6 motor root bursting for all illumination durations: 10s (pre: 9.6±7.5 m^-1^, on: 55.4±26.1 m^-1^, p = .002), 20s (pre: 11.4±4.6 m^-1^, on: 46.3±9.1 m^-1^, p=.002), and 40s (pre: 11.2±6.6 m^-1^, on: 41.0±12.9 m^-1^, p=.005) (Figure 4A, Ci) and a decrease in bursting duration for all illumination durations: 10s (pre: 2.8±1.3 m^-1^, on: 1.0±0.4 m^-1^, p=.002) 20s (pre: 3.3±1.2 m^-1^, on: 0.9±0.3 m^-1^, p=.004), and 40s (pre: 3.2±1.7 m^-1^, on: 1.7±1.9 m^-1^, p=.005) stimulation periods (Figure 4Cii). Elevated A6 motor root bursting frequency was also evoked by pulsed-based stimulation (Supplementary Figure 1). Across both continuous and pulsed-based stimulation of *tdc*^*2+*^ neurons, intrinsic manipulation of the *tdc*^*2+*^ system induced fictive forward bias akin to results demonstrated by exogenous OA application (Figure 4E). Specifically, optogenetic depolarisation of *tdc*^*2+*^ neurons induced increased fictive forward motor bias for all illumination durations: 10s (pre: 3.3±2.2 m^-1^, on: 64.2±43.0 m^-1^, p < .001), 20s (pre: 4.3±4.0 m^-1^, on: 36.3±7.7 m^-1^, p < .001), and 40s (pre: 4.8±5.1 m^-1^, on: 36.8±10.8 m^-1^, p <.001) Across stimulation periods, optogenetic depolarisation of *tdc*^*2+*^ neurons induced fictive forward waves in 79±8% of all fictive activity events during stimulation from 36±13% before and 36±14% after stimulation.

**Figure 4:**
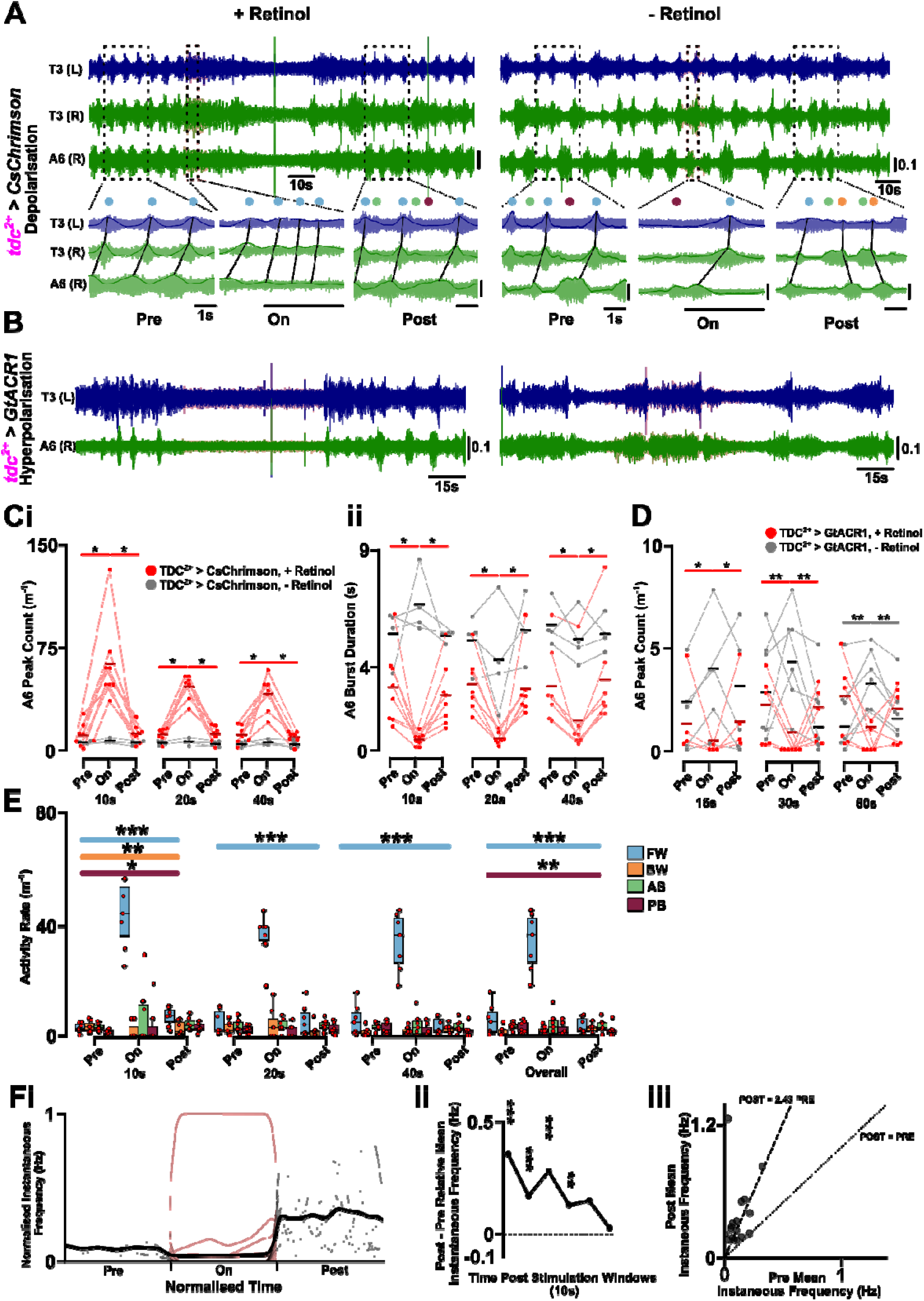
*Tdc*^*2+*^ Neuronal Activity is Sufficient and Necessary for Fictive Locomotor Activity. (**A**) Exemplar three nerve root (T3 (L), T3 (R), A6(R)) electrophysiology trace for continuous red-light depolarisation of *tdc*^*2+*^ neurons using CsChrimson for preparations raised on retinol (experimental, left) and no retinol (control, right). Before, during (red box), and post-stimulation periods are shown with rectified and smoothed traces shown. Black lines represent fictive activity motifs. (B) Exemplar two nerve root (T3 (L), A6 (R)) electrophysiology trace for continuous white-light hyperpolarisation (red box) of *tdc*^*2+*^ neurons using GtACR1 for preparations raised on retinol (experimental, left), and no retinol (control, right). (C) Quantification of abdominal (A6) motor root bursting for (i) burst rate and (ii) duration of bursts in *tdc*^*2+*^ >CsChrimson for 10s, 20s, and 40s depolarisation period. (D) Quantification of abdominal (A6) motor root bursting for burst rate in *tdc*^*2+*^ >GtACR1 for 15s, 30s, and 60s hyperpolarisation period. (E) Change in the fictive activity profile for *tdc*^*2+*^ >CsChrimson stimulation periods demonstrating inducing of fictive forward bias across stimulation period. Activity profile within stimulation periods and average across stimulation periods are shown. (Fi) Changes in normalised instantaneous frequency of overall fictive activity motor patterns before, during, or after hyperpolarisation of *tdc*^*2+*^ neurons (N=30 repeats over N=7). (Fii) Relative change in fictive frequency instantaneous frequency in 10 second intervals after optogenetic hyperpolarisation compared to pre-stimulation overall fictive activity instantaneous frequency. (Fiii) Mean instantaneous frequency of post-inhibition and pre-inhibition period demonstrating a significant post-inhibitory rebound in fictive activity within 1-minute post-inhibition. Comparisons followed by post-hoc statistics were conducted with p-values: < .5 (*), < .01 (**), <.001 (***) shown. Only statistically significant bars are show

Secondly, we optogenetically hyperpolarised *tdc*^*2+*^ neurons expressing GtACR1 with pulses of white light (15s, 30s, 60s) and recorded motor root activity from left and right T3 and A6 VNC ganglia (Figure 4B). Optogenetic hyperpolarisation of *tdc*^*2+*^ neurons reduced, and in many cases, completely abolished motor root bursting for all light-stimulation periods: 15s (pre: 1.6±2.1 m^-1^, on: 0.5±1 m^-1^, p = .04), 30s (pre: 1.8±1.8 m^-1^, on: 0±0 m^-1^, p=.009), and 60s (pre: 2.2±2.1 m^-1^, on: 0.4±0.7 m^-1^, p = .01) illumination periods with 37 of 49 (75.9±30.8%) trials across preparations (N=7) showing no fictive activity during the illumination period (Figure 4Bii, D). For longer illumination periods (> 15s), there was weak recovery of rhythmic activity in the A6. Interestingly, long-lasting (60s) LED white-light illumination in *tdc*^*2+*^ *>GtACR1* control preparations, in which the optogenetic tools in activated by withholding the essential cofactor retinol, induced significantly increased (pre: 1.8±2.0 m^-1^, on: 3.7±1.5 m^-1^, p = .01; Figure 4D) A6 motor root activity potentially indicative of photoreceptor-based motor stimulation in the isolated VNC preparation.

Optogenetic inhibition of *tdc*^*2+*^ neurons induces post-inhibitory rebound in fictive activity (Figure 4F). By normalising the instantaneous frequency of all fictive activity across inhibition period windows (1-minute pre, on, post), optogenetic inhibition of *tdc*^*2+*^ neurons suppresses activity and induces an elevation in the rate of fictive activity immediately after the cessation of inhibition (Figure 4Fi). Through comparing the post-stimulation to pre-stimulation instantaneous frequency of all fictive activity, there is a significant elevation in activity rate within 0-40s after removal of optogenetic inhibition (0-10s: 0.35±0.12Hz, p < .001; 10-20s: 0.17±0.04Hz, p < .001; 20-30s: 0.28±0.12Hz, p < .001; 30-40s: 0.13±0.04Hz, p = .002) (Figure 4Fii). After optogenetic inhibition, the mean instantaneous frequency of fictive activity assessed via regression-through-origin was 2.43±1.94 times that of the pre-stimulation period (Figure 4Fiii). Taken together, inhibiting *tdc*^*2+*^ neuron activity collapses fictive rhythms and reveals a post-inhibitory rebound in the fictive landscape for 40s beyond manipulation.

Overall, *tdc*^*2+*^ neuron depolarisation increased fictive forward activity akin to exogenous OA application (see accompanying paper) and hyperpolarisation of these neurons collapsed all motor rhythms. Thus, these results demonstrate *tdc*^*2+*^ neurons have command-like properties for fictive motor activity in isolated VNC 3^rd^ *Drosophila* larval preparations.

Given the *tdc*^*2+*^ neuron population consists of local and projecting neurons, we next turned to spatially-restricting depolarisation to assess the contributions of subpopulations of VNC octopaminergic neurons. We restricted red-light illumination of *tdc*^*2+*^ neuronal populations in the ventromedial axis of the isolated ventral nerve cord using a custom-built physiology set-up. Red-light illumination of *tdc*^*2+*^ neuronal populations increased bursting rate in all restricted regions: SOG (Figure 5A), thoracic ganglion (Figure 5B), anterior abdominal ganglion (Figure 5C), and posterior abdominal ganglion (Figure 5D). Across all stimulation regions, both the thoracic and abdominal nerve root bursting rate is increased during *tdc*^*2+*^ depolarisation (Figure 6Ai - Di): illuminating the SOG induced a increase in T3 (pre: 8.3±4.8 m^-1^, on: 30.0±6.9 m^-1^, p = .001) and A6 (pre: 8.8±4.7 m^-1^, on: 31.7±10.8 m^-1^, p = .001) (Figure 6Ai); illuminating thoracic ganglion (T1-3) induced a increase in T3 (pre: 6.5±3.4 m^-1^, on: 30.0±6.0 m^-1^, p < .001) and A6 (pre: 6.8±3.5 m^-1^, on: 37.7±13.3 m^-1^, p < .001) (Figure 6Bi); illuminating anterior abdominal ganglion (A1-3) induced a increase in T3 (pre: 6.6±2.1 m^-1^, on: 34.3±12.4 m^-1^, p = .001) and A6 (pre: 6.8±3.5 m^-1^, on: 32.6±17.3 m^-1^, p = .022) (Figure 6Ci); and illuminating posterior abdominal ganglion (A6-8) induced a increase in T3 (pre: 6.1±2.5 m^-1^, on: 48.0±12.0 m^-1^, p < .001) and A6 (pre: 6.5±2.1 m^-1^, on: 45.4±9.7 m^-1^, p < .001) (Figure 6Di). Further, across all stimulation regions, both the thoracic and abdominal nerve root bursting duration was significantly reduced during *tdc*^*2+*^ depolarisation with the exception of the T3 motor root during thoracic and anterior abdominal ganglion (Figure 6Ai-Di). Specifically, direct SOG *tdc*^*2+*^ depolarisation induced a significant decrease in T3 (pre: 3.0±1.3s, on: 1.5±0.6s, p = .01) and A6 (pre: 3.0±1.3s, on: 1.3±0.4s, p = .03) motor root burst duration (Figure 6Ai). While direct thoracic ganglion (T1-3) *tdc*^*2+*^ depolarisation did not induced a significant decrease in T3 motor root burst rate (pre: 2.9±1.0s, on: 1.5±0.4s, p = n.s), depolarisation of T1-T3 *tdc*^*2+*^ neurons did induce a significant decrease in the A6 motor root burst duration (pre: 3.1±1.9s, on: 1.4±0.6s, p = .02) (Figure 6Bi). While direct anterior abdominal ganglion (A1-3) *tdc*^*2+*^ depolarisation did not induced a significant decrease in T3 motor root burst rate (pre: 3.2±1.2s, on: 1.5±0.4 m^-1^, p = n.s), depolarisation of A1-A3 *tdc*^*2+*^ neuron did induce a significant decrease in the A6 motor root burst duration (pre: 3.1±1.9s, on: 1.4±0.6s, p = .04) (Figure 6Ci). Finally, direct posterior abdominal ganglion (A6-8) *tdc*^*2+*^ depolarisation induced a significant decrease in T3 (pre: 3.5±1.3s, on: 1.1±0.2s, p = .001) and A6 (pre: 3.3±1.1s, on: 1.1±0.2s, p = .003) motor root burst duration (Figure 6Di). Thus, depolarisation of *tdc*^*2+*^ neuron subpopulations in VNC ganglia is sufficient to induce elevated motor activity in the isolated ventral nerve root.

**Figure 5:**
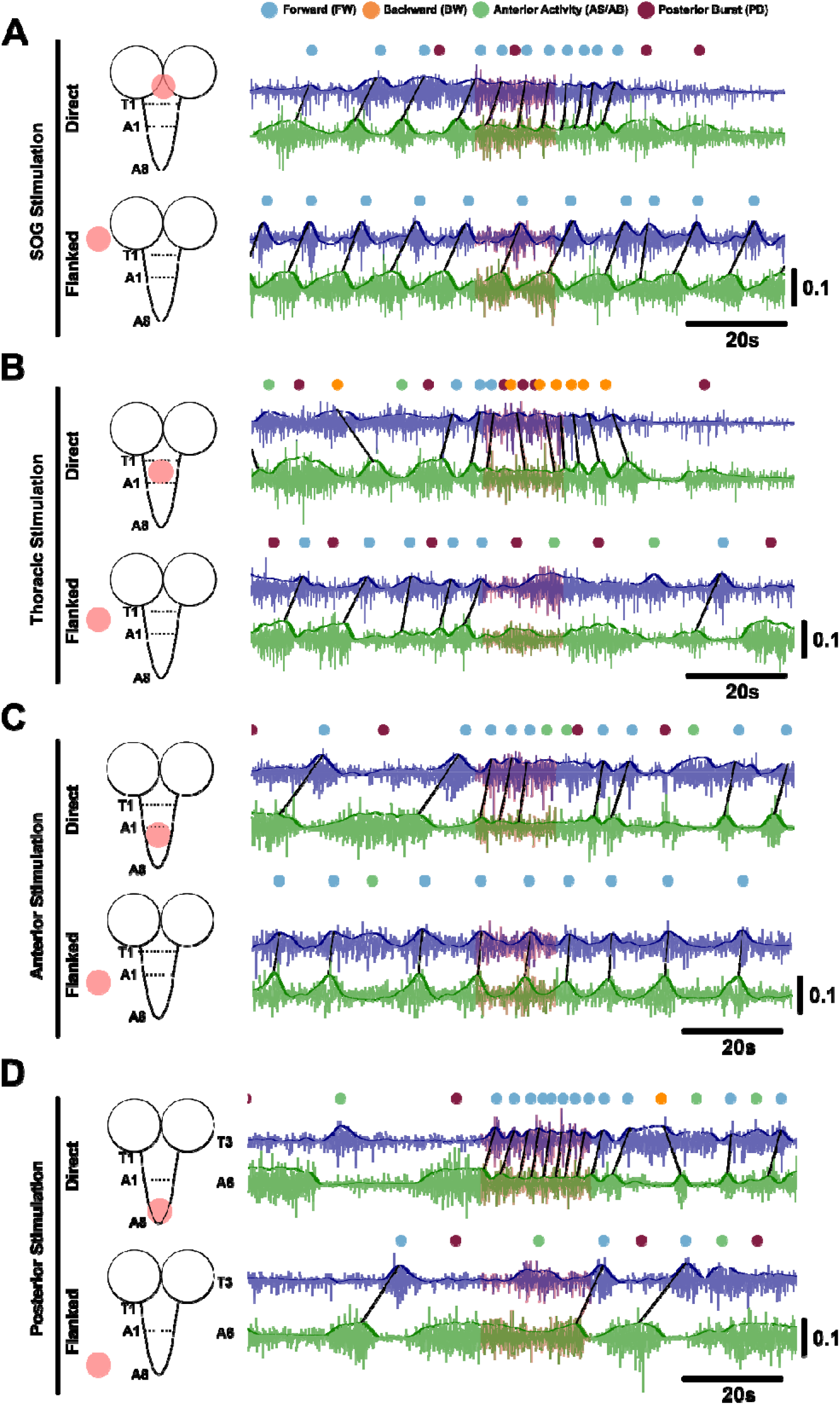
Restricted Optogenetic Stimulation of *tdc*^*2+*^ Neurons Induces Diversity of Activity. Restricted 10s optogenetic stimulation of *tdc^2+^* neurons in the (**A**) SOG, (**B**) thoracic ganglion (T1-T3), (**C**) anterior abdominal ganglion (A1-A3), and (**D**) posterior abdominal ganglion (A6-A8). Representative electrophysiology thoracic (T3) and posterior abdominal (A6) nerve root recording traces for direct and flanked stimulation conditions shown with coloured-dots indicating occurrence of fictive motor pattern: fictive forward wave (blue dot), posterior burst (purple dot), fictive backward wave (orange dot), anterior a/symmetric activity (green dot).

We next examined if restricted *tdc*^*2+*^ neuron depolarisation induced fictive forward bias akin to exogenous OA application. Depolarisation of *tdc*^*2+*^ neurons in the SOG (Figure 6Aii), thoracic ganglion (Figure 6Bii) and anterior abdominal ganglion (Figure 6Cii) did not clearly bias fictive behavioural patterns. However, individual iterations of restricted stimulation across preparations did demonstrate a significant variation in the type of fictive bias within each region of *tdc*^*2+*^ depolarisation (Figure 5, 6). Namely, restricted *tdc*^*2+*^ depolarisation in the SOG (Figure 6A, χ^2^(2) = 7.3, p = .007), thoracic ganglion (Figure 6B, χ^2^(2) = 7.3, p = .007), and anterior abdominal ganglion (Figure 6C, χ^2^(2) = 7.3, p = .007) all significant induced changes in fictive wave bias with some instances showing promotion of fictive forward whereas others fictive backward bias. In contrast, posterior abdominal restricted *tdc*^*2+*^ depolarisation induced a significant increase in fictive forward motor activity (pre: 2.4±1.9m^-1^, on: 38.6±13.8m^-1^, p = .005) (Figure 6Dii). During posterior abdominal restricted *tdc*^*2+*^ depolarisation a majority of stimulation instances were fictive forward biased (Figure 6D, χ^2^(2) = 5.8, p = .02). Thus, depolarising abdominal posterior *tdc*^*2+*^ neurons, with or without depolarising all VNC *tdc*^*2+*^ neurons, induces a bias towards fictive forward motor bias.

**Figure 6:**
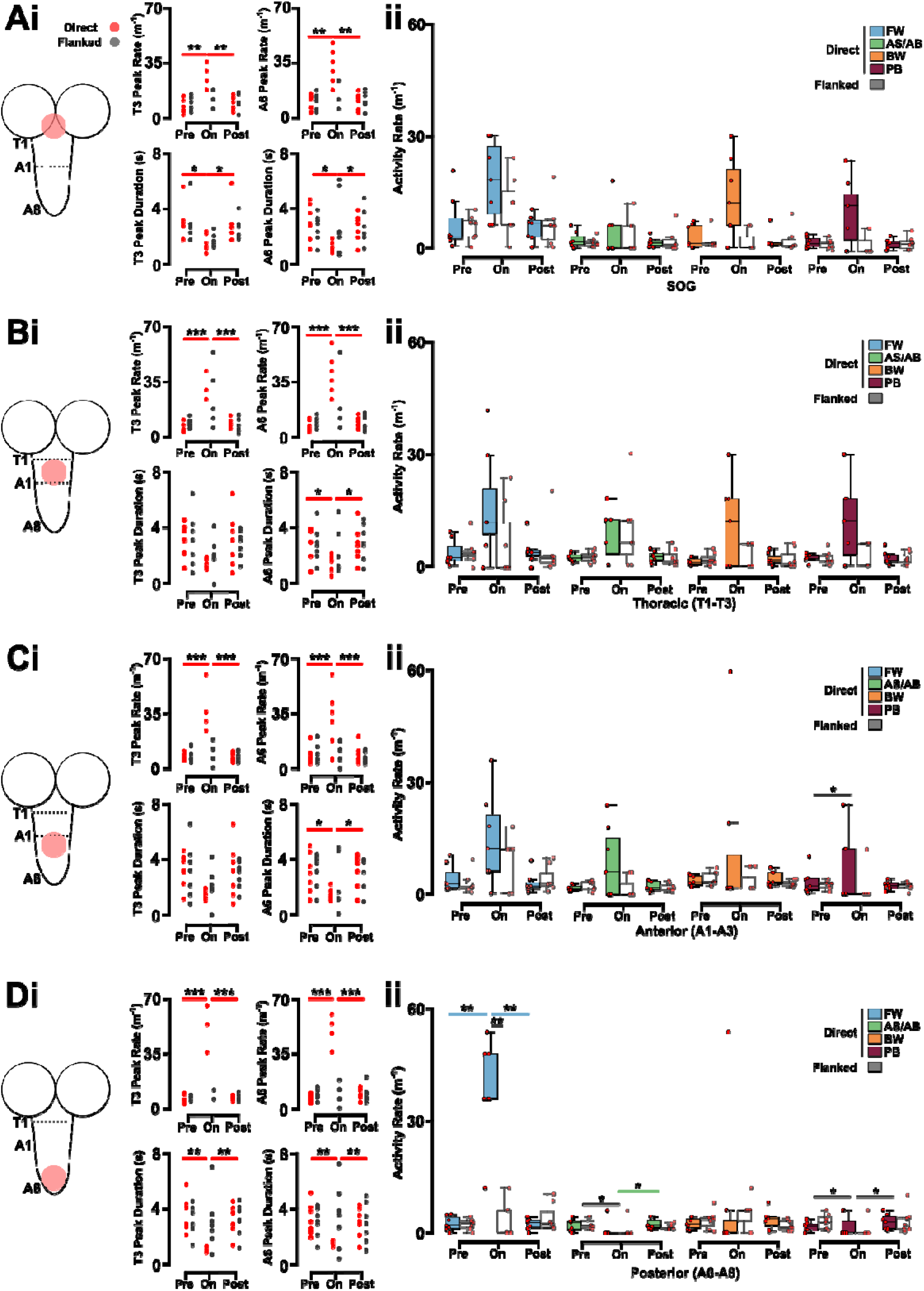
Optogenetic Stimulation of Posterior Abdominal *tdc*^*2+*^ Neurons Induces Fictive Forward Locomotion. Quantification of (i) abdominal and thoracic burst rate, burst duration, and (ii) fictive activity (“FW” = fictive forward wave, “AS/AB” = anterior symmetric or asymmetric, “BW” = fictive backward wave, “PB” = posterior bust) rate during direct (colour) and flanked (grey) optogenetic stimulation of (A) SOG, (B) thoracic ganglion (T1-T3), (C) anterior abdominal ganglion (A1-A3), and (D) posterior abdominal ganglion (A6-A8) *tdc*^*2+*^ neurons. Comparisons followed by post-hoc statistics were conducted with p-values: < .5 (*), < .01 (**), <.001 (***) shown. Only statistically significant bars are shown.

### Descending Tdc^2+^ Neurons are Uniquely Responsible for Optogenetic-induced Bias in Fictive Activity

Given there exist both local and descending *tdc*^*2+*^ neurons, we turned to using restricted expression to isolate which population of *tdc*^*2+*^ neurons are uniquely inducing the promotion of fictive activity. First, we confirmed the restricted expression of *CsChrimson* to either the descending brain *tdc*^*2+*^ neurons (Supplementary Figure 2) or exclusively the VNC-residing *tdc*^*2+*^ neurons using the *tsh*^*2+*^ construct (Supplementary Figure 3). Qualitatively, optogenetic stimulation of exclusively VNC-residing *tdc*^*2+*^ neurons (N=5) did not induce a change in fictive bias (Figure 7A) whereas stimulation of brain-residing *tdc*^*2+*^ neurons (N=7) was sufficient to induce fictive forward bias (Figure 7B). Specifically, the fictive forward promotive bias was inducible by posterior-restricted stimulation of brain-residing *tdc*^*2+*^ neurons (Figure 7B, posterior stimulation). Optogenetic stimulation of VNC-residing *tdc*^*2+*^ neurons did not induce any significant increases in bursting, decreases in bursting amplitude (Figure 8Ai), nor any increases in fictive activity (Figure 8Aii). In contrast, VNC-wide optogenetic stimulation of brain-residing *tdc*^*2+*^ neurons induced a significant increase in bursting frequency in both the thoracic (pre: 5.8±2.5m^*-1*^, on: 21.0±14.2m^-1^, p = .006) and abdominal segments (pre: 5.3±3.6m^-1^, on: 17.3±7.7m^-1^, p = .004) while significantly decreasing the bursting duration of abdominal segments (pre: 8.7±10.7s, on: 3.4±1.0s, p = .01) (Figure 8Bi). Further, VNC-wide optogenetic stimulation of brain-residing *tdc*^*2+*^ neurons induce a significant increase in uniquely fictive forward activity (pre: 2.7±1.6m^-1^, on: 25.2±14.4m^-1^, p = .004) alongside a significant reduction in posterior burst activity (pre: 1.3±0.9m^-1^, on: 0.3±0.5m^-1^, p = .028) (Figure 8Bii). Restricted stimulation of the brain-residing burst activity *tdc*^*2+*^ neuron’s posterior-extending neuropils similarly induced a significant increase in bursting frequency in the abdominal segments (pre: 4.8±2.6m^-1^, on: 19.9±10.7m^-1^, p = .03) while significantly decreasing the bursting duration of abdominal segments (pre: 4.9±1.6s, on: 2.5±1.6s, p = .03) (Figure 8Ci). Further, VNC-wide optogenetic stimulation of brain-residing *tdc*^*2+*^ neurons induce a significant increase in uniquely fictive forward activity (pre: 1.5±1.0m^-1^, on: 22.2±7.8m^-1^, p = .009) and significant reduction in fictive backward activity (pre: 1.6±1.4m^-1^, on: 0±0m^-1^, p = .014) (Figure 8Cii).

**Figure 7:**
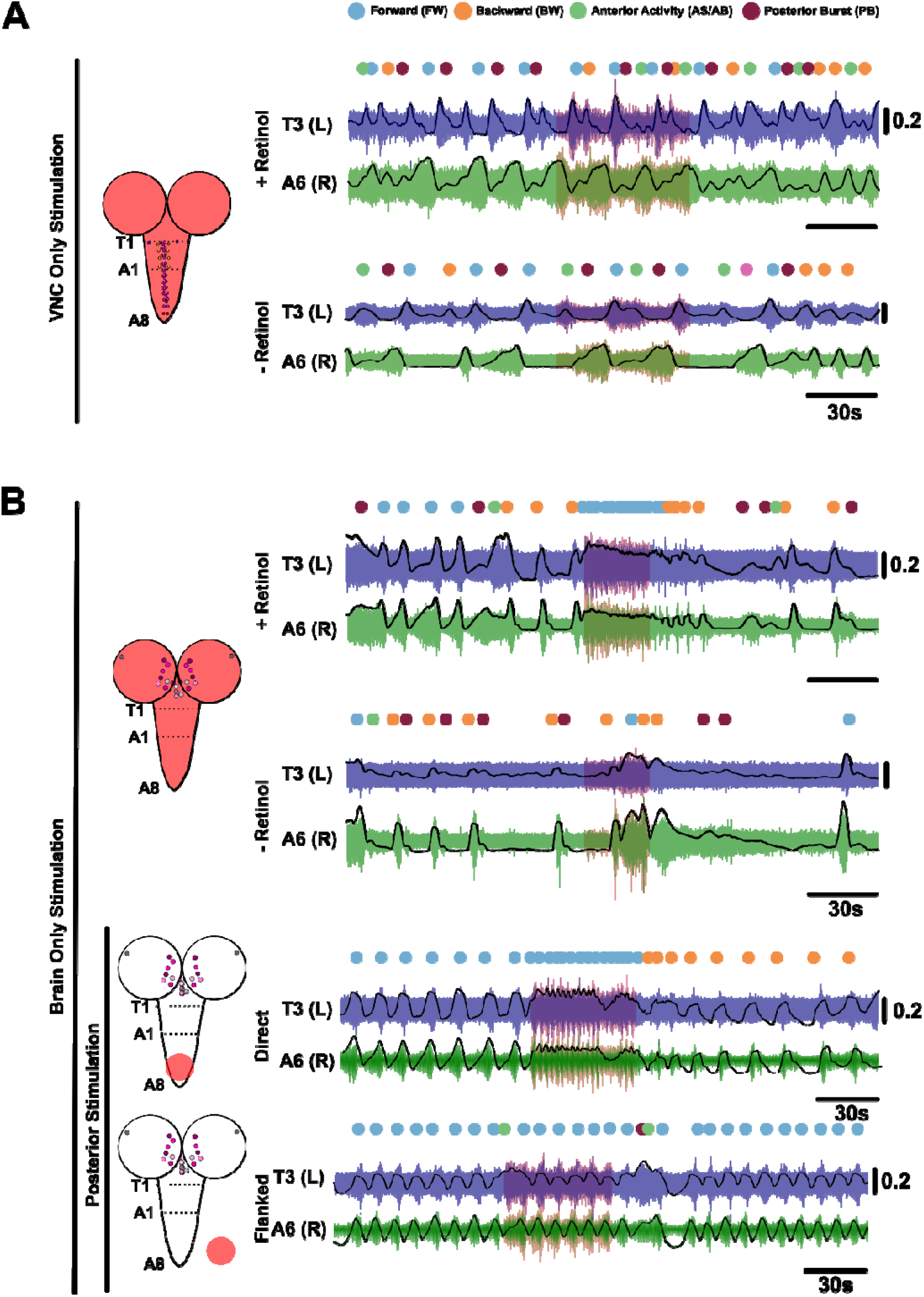
Optogenetic Stimulation of Descending *tdc*^*2+*^ Neurons Recapitulate Bias to Fictive Forward Activity. (**A**) Neve root activity from system-wide optogenetic stimulation of VNC-residing *tdc*^*2+*^ neurons within *Drosophila* reared either retinol (N=5) or no retinol (control) (N=3). (**B**) Neve root activity from system-wide optogenetic stimulation of brain-residing *tdc*^*2+*^ neurons within *Drosophila* reared either retinol (N=7) or no retinol (control) (N=3). Restricted light on the posterior segments (A6-8) stimulating posterior-projected brain-residing *tdc*^*2+*^ neurons neuropils (N=7) and flanked control (N=4). Fictive activity patterns are annotated above traces: fictive forward wave (blue dot), posterior burst (purple dot), fictive backward wave (orange dot), anterior a/symmetric activity (green dot).

**Figure 8:**
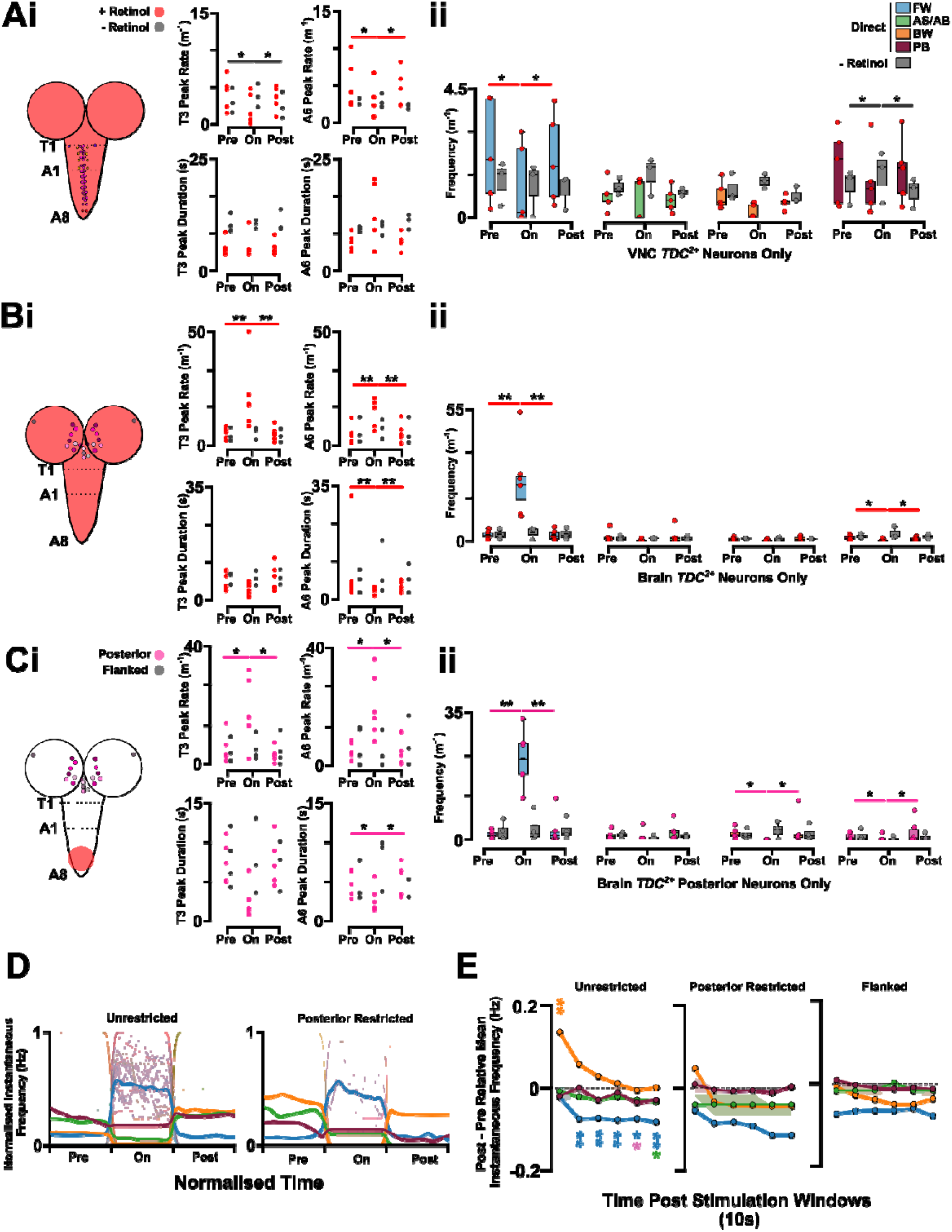
Optogenetic Stimulation of Descending *tdc*^*2+*^ Neurons Induce a Transient Increase in Fictive Backward Activity. (**A**) The effect of optogenetic stimulation of system-wide VNC-residing *tdc*^*2+*^ neurons on (**i**) thoracic and abdominal motor root bursting frequency and bursting duration alongside (**ii**) changes in the frequency of fictive activities. (**B**) The effect of optogenetic stimulation of system-wide brain-residing *tdc*^*2+*^ neurons on (**i**) thoracic and abdominal motor root bursting frequency and bursting duration alongside (**ii**) changes in the frequency of fictive activities. (**C**) The effect of optogenetic stimulation of posterior-restricted brain-residing *tdc*^*2+*^ neurons on (**i**) thoracic and abdominal motor root bursting frequency and bursting duration alongside (**ii**) changes in the frequency of fictive activities. (**D**) Changes in normalised instantaneous frequency of fictive motor patterns as a result of unrestricted, system-wide (left) and posteriorly-restricted (right) optogenetic stimulation of brain-residing *tdc*^*2+*^ neurons. (**E**) Relative change in fictive frequency instantaneous frequency in 10 second intervals after optogenetic stimulation compared to pre-stimulation fictive activity instantaneous frequency. Comparisons followed by post-hoc statistics were conducted with p-values: < .5 (*), < .01 (**), <.001 (***) shown.

In an accompanying paper, we show that octopaminergic neuromodulation via perfusion onto isolated *Drosophila* larval ventral nerve cords can induce post-hoc changes in the dominant fictive profile after application (i.e., anterior asymmetry promotion after the removal of octopaminergic-induced fictive forward bias) [33]. There was a significant immediate elevation in the immediate instantaneous frequency for fictive backward activity during system-wide descending stimulation of *tdc*^*2+*^ neurons (relative increase of .14Hz in 0-10s post stimulation: p =.007, N = 31) (Figure 7D, E). Additionally, there was a longer-sustained significant reduction in the instantaneous frequency of fictive forward activity post fictive induction via *tdc*^*2+*^ neurons stimulation (relative reduction of .07±.15Hz in 0-60s post stimulation period: p = .003, N=56). Overall, optogenetic stimulation of descending *tdc*^*2+*^ neurons – plausibly sVPMmd1, sVUMlb3, tyraminergic ICA neurons [8] – were sufficient to induce the promotion of fictive forward activity with sustained motor bias inducing reactive, prolonged changes in the fictive motor landscape of the isolated *Drosophila* preparation.

## Discussion

The adrenergic-like system appears associated with the initiation and modulation of a diversity of behaviours across invertebrates. Here, we show that the adrenergic-like *tdc*^*2+*^ system tracks fictive activity and plays a critical role in generating rhythmic activity in *Drosophila* larval CPG networks. Specifically, we show that stimulation of descending *tdc*^*2+*^ neurons – plausibly octopaminergic sVPMmd1 and sVUMlb3 or tyraminergic ICA neurons – appears sufficient to induce fictive forward rhythms reminiscent of activity during octopamine bath application. We also demonstrate that hyperpolarizing *tdc*^*2+*^ neurons collapses rhythmogenesis in the larval CNS, suggesting that they may be generally necessary for rhythmogenesis in this system.

### The Role of *tdc*^*2+*^ Activity in Drosophila Larvae Fictive Activity

Previous work has characterised the importance of *tdc*^*2+*^ neurons for coordinated intact locomotion [12], here we demonstrate VNC-residing *tdc*^*2+*^ neurons preceded all fictive motor activity, brain-descending *tdc*^*2+*^ neurons are sufficient for fictive forward program initiation, and inhibition of *tdc*^*2+*^ neurons collapses all motor rhythms. The phase locking of VNC-residing *tdc*^*2+*^ neurons activity to motor patterns is unsurprising considering the general role of adrenergic-like modulation in shaping motor activity non-selectively. Several *Drosophila* locomotor circuits contain neurons whose activity tracks or precede motor output (i.e., A18b/Pair1 in larval backwards activity [34], AcN/A01j/A02j circuits in fast forward locomotion [35], and adult descending neurons that predict walking or steering [36]). Thus, *tdc*^*2+*^ neurons activity preceding multiple fictive programs extends this principle to the adrenergic-like system implying *tdc*^*2+*^ neurons may provide a broad motor-readiness or gain-setting signal upstream of programme-specific premotor circuits.

Restricted optogenetic stimulation of the most posterior ganglion segments (A6-8) was reliably able to induce fictive forward locomotion. As shown in cell-level staining [8], [12], restricting expression to *tdc*^*2+*^ neurons in the brain results in expression in both descending *tdc*^*2+*^ neurons and local *tdc*^*2+*^ neurons in the aDM9 cluster. Specifically, only 3 sets of brain-residing *tdc*^*2+*^ neurons project to the segments we optogenetically stimulated – sVPMmd1, sVUMlb3, ICA [8]. Given optogenetic stimulation of exclusively VNC-residing *tdc*^*2+*^ neurons was unable to induce fictive forward bias, we can infer the three set of descending *tdc*^*2+*^ neurons may play a significant role in the reported rhythmogenetic bias properties. Given SOG stimulation plausible depolarises sVPMmd1-3 and sVUMlb1-3 whereas posterior VNC stimulation plausible depolarises exclusively sVPMmd1 and sVUMlb3 neurons and SOG stimulation induces a diverse array of different fictive motor programs whereas posterior stimulation induces exclusively fictive forward activity, distinct parts of the *tdc*^*2+*^ system may play distinct roles in facilitating distinct motor programs. Within adult *Drosophila* rhythmogenesis, descending *tdc*^*2+*^ neurons projecting to the thoracic ganglion play an individual and unique role in locomotion [37]. Indeed, more broadly VUM/DUM neurons across insects are characteristically differentially recruited and modulate motor states (e.g., flight CPG) [12], [17], [38], [39]. Beyond rhythmogenesis, sVUMlb3 may operate as a sensory integrator shaping behavioural computations like olfactory discrimination [15] with VPM *tdc*^*2+*^ neurons having broader roles in shaping behavioural-state bias via mushroom-body modulation (e.g., appetitive learning [17], [40], [41], arousal and sleep [42], [43]). While our results imply a command-like property, to establish and ascribe a direct, command-like rhythmogenic property to sVPMmd1, sVUMlb3, or ICA future work may utilise stochastic or split-GAL4 with excitatory and inhibitory optogenetics to produce a functional map of individual *tdc*^*2+*^ neuron influence on fictive activity.

### Constraints for a *tdc*^*2+*^ -Motor System Model

While the cellular architecture of adrenergic-like *tdc*^*2+*^ system in *Drosophila* has been reported [8] [44], how essential this system is for core rhythmogenesis is underexplored. In addition to our exogenous modulation work [33], we evidence key features of rhythm generation that could be used to constrain future ensemble and/or connectome-based modelling of motor system architecture. The functional properties of *tdc*^*2+*^ neurons reported here concurs with prior anatomical assessment providing a strong basis for future modelling architecture. VNC-residing *tdc*^*2+*^ neurons appear to exclusively project into the body-wall musculature [8], [32], [45], [46], [47]. Indeed, a significant level of octopaminergic modulation is exhibited at the neuromuscular junction via the VNC-residing *tdc*^*2+*^ neurons [9], [48]. Given *tdc*^*2+*^ neuron activity was highly coupled to all fictive activity patterns, *tdc*^*2+*^ activity may represent a broad motor-readiness or gain-setting signal rather than a behaviour-specific motor command. However, given that optogenetic stimulation of *tdc*^*2+*^ neuron induced predominately fictive forward activity rather than generalized increases in motor activity, it is clear that the *tdc*^*2+*^ system occupies a permissive and biasing position in the motor hierarchy underlying rhythm generation. Previously, we have shown how motor programme diversity and variability can arise from interactions amongst inhibitory circuit motifs embedded in the segmentally-organised larval locomotor system [3]. Given that stimulation of descending *tdc*^*2+*^ neurons does not just induce fictive forward behaviour, but significantly quicker waves in more intense bouts, the *tdc*^*2+*^ neurons may be generally synaptically connected to not just command-like structures for distinct patterns, but also the interneuron structures responsible for their intersegmental propagation and thus progression speed. Taken together, the *tdc*^*2+*^ system likely operates upstream of putative inhibitory motifs and interneuron patterning circuits. Thus, *tdc*^*2+*^ neurons activity may alter the gain, threshold, or recovery dynamics of inhibitory motif modules, thereby providing the potential to change which motor program dominates the network without eliminating the capacity of the network to generate diverse outputs. Conceptually, this work generates several testable predictions for any modelled *tdc*^*2+*^ component: increased the *tdc*^*2+*^ activity should reduce the threshold for fictive forward wave initiation, particularly for posterior-projecting components; in turn, *tdc*^*2+*^ neuron inhibition should reduce the probability of all fictive programmes initiating by lowering the excitability of rhythmogenetic modules or preventing inhibitory modules from escaping inactivity.

### The Adrenergic-like System in Modulating Motor Competition

Sustained application of octopamine to the *Drosophila* larval preparation induces prolonged fictive forward motor bias with a sustained elevation in anterior asymmetric activity post application (accompanying paper, [33]). Responsivity in any dynamic system is one that is reliably sensitive to changing exogenous and endogenous demands and changes only when appropriate. In the context of neural systems, particularly motor systems, a diverse array of potential outputs is possible – different patterns (e.g., forwards vs backwards), different speeds, different sequences of activity. Thus, a motor system must maintain the capacity to be sensitive so it can alter its output when appropriate, but also remain resilient against inappropriate, maladaptive activity patterns. Here, we show that optogenetic manipulation of the *tdc*^*2+*^ system can induce a reversible change in the fictive landscape of activity – but also promotes a competing motor program (backward waves) after cessation of stimulation. It remains to be determined if the reactive changes in the fictive landscape are a passive, or active, component in recovering the motor network to a state of normal output diversity. Conceptually, it may be the case *tdc*^*2+*^ neurons exhaust forward-specific components of the network, thus a prolonged refractory period emerges. However, such a refractory period would need to extend into the realm of minutes, plausibly eliminating many cell-level refractory mechanisms. Thus, synaptic-level or circuit-level mechanism are plausibly responsible for the alternations in the fictive landscape post bias induction. Similarly, given *tdc*^*2+*^ neuron suppression evokes elevated post-inhibition rhythmicity, it implies that the *tdc*^*2+*^ system is simultaneously linked to general rhythm generation, but is capable of engagement the forward circuit independently under certain conditions. Given that previous work has shown the *Drosophila* larval VNC most likely contains independent oscillators in each hemi-segment, it may be the case the *tdc*^*2+*^ system synapses, directly or indirectly, to hemisegmental oscillators at distal ends of the network which are responsible for triggering cascading waves across coupled oscillators [49]. Future work can assess this by optogenetically manipulating specific *tdc*^*2+*^ neurons that may project to partners in the thoracic ganglion (i.e., for backward/asymmetry generation [3]) or other subcomponents of the inhibitory networks in distal regions [3], [49]. Overall, this work – alongside our prior work [33] and an accompanying paper – demonstrates core principles, circuit components and functional constraints for monitoring and reinforcing diversity amongst competing central pattern generating networks.

## Conflicts of Interest

No declared conflict of interest.

## Data Availability

The research data underpinning this publication can be accessed at https://doi.org/10.17630/98cb0f9f-f064-4d08-addf-51d0025d1a7c.

## Author’s Contribution

S.R.P. and W.V.S. conceived the study. W.V.S conducted all experiments, analysis, and visualisation. W.V.S. and S.R.P finalised the figures. S.R.P. supervised the project.

## Acknowledgements

This project was made possible by an Industrial CASE PhD studentship (UKRI Biotechnology and Biological Sciences Research Council (BBSRC) grant number BB/T00875X/1) to WVS. This work was additionally supported in part by two Collaborative Research Grants awarded jointly by the Global Office of the University of St Andrews and The Halle Institute for Global Research at Emory University (SRP and A. Prinz), and The Global Office of the University of St Andrews and the University of Bonn (SRP and M. Pankratz).

## Supplementary Figures

**Supplementary Figure 1:**
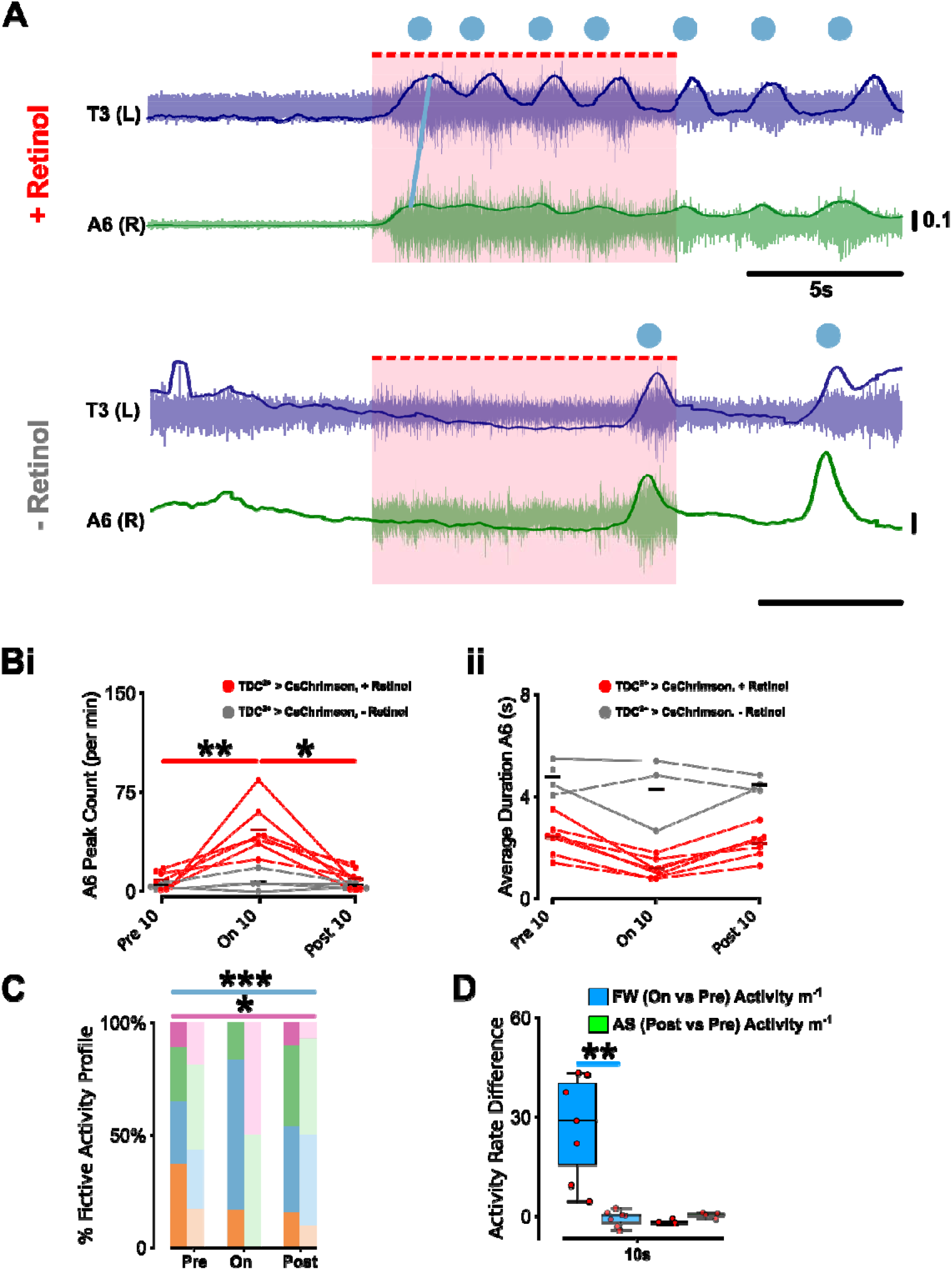
Pulsed Stimulation of *tdc*^*2+*^ Neurons Induces Fictive Forward Bias. (**A**) Representative T3(L) and A6(R) motor root electrophysiology trace demonstrating promotion of fictive forward activity during pulsed stimulation of CsChrimson>*tdc*^*2+*^ (0.01s, 40Hz, 400 repeats = 10s total stimulation) for experimental (+ retinol) compared to control (-retinol) 3^rd^ instar *Drosophila* larva. (**B**) Quantification of motor root bursting at A6 demonstrating (**i**) peak rate and (**ii**) burst duration changes upon pulsed stimulation (On10) relative to before (Pre10) and after (Post10) stimulation period. (**C**) Quantification of fictive bias before, during, and after pulsed stimulation in preparations where fictive activity could be accurately inferred (N=6). (**D**) The relative fictive activity difference for fictive forward activity during (on) vs before (pre) stimulation (blue) and for anterior asymmetric fictive activity after (post) vs before (pre) stimulation (green). Comparisons followed by post-hoc was conducted with p-values: < .5 (*), < .01 (**), <.001 (***) shown. Only statistically significant bars are shown.

**Supplementary Figure 2:**
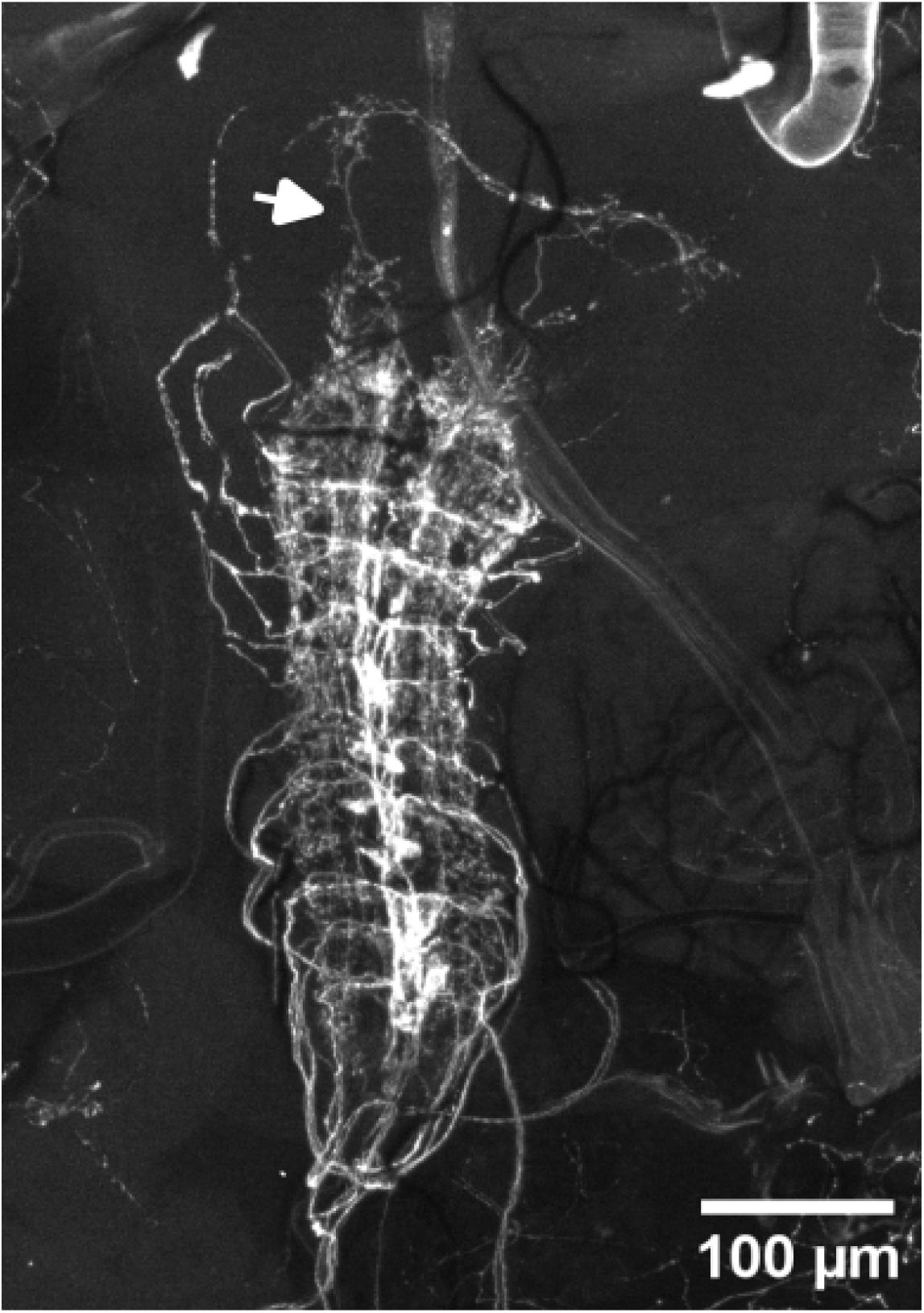
Restricted *tdc*^*2+*^ Neuron Anatomy to the VNC. Confocal staining of 20XUAS>dsFRT>CsChrimson, tsh-LexA, 8xLexAopFlp attP40/CyO, Tb-RFP with w[*]; P{w[±mC]=Tdc2-GAL4 at 10x magnification. Note the absence of *tdc*^*2+*^ neurons soma in the brain lobes. Ascending projections from the VNC to the brain lobe are shown via a white arrow.

**Supplementary Figure 3:**
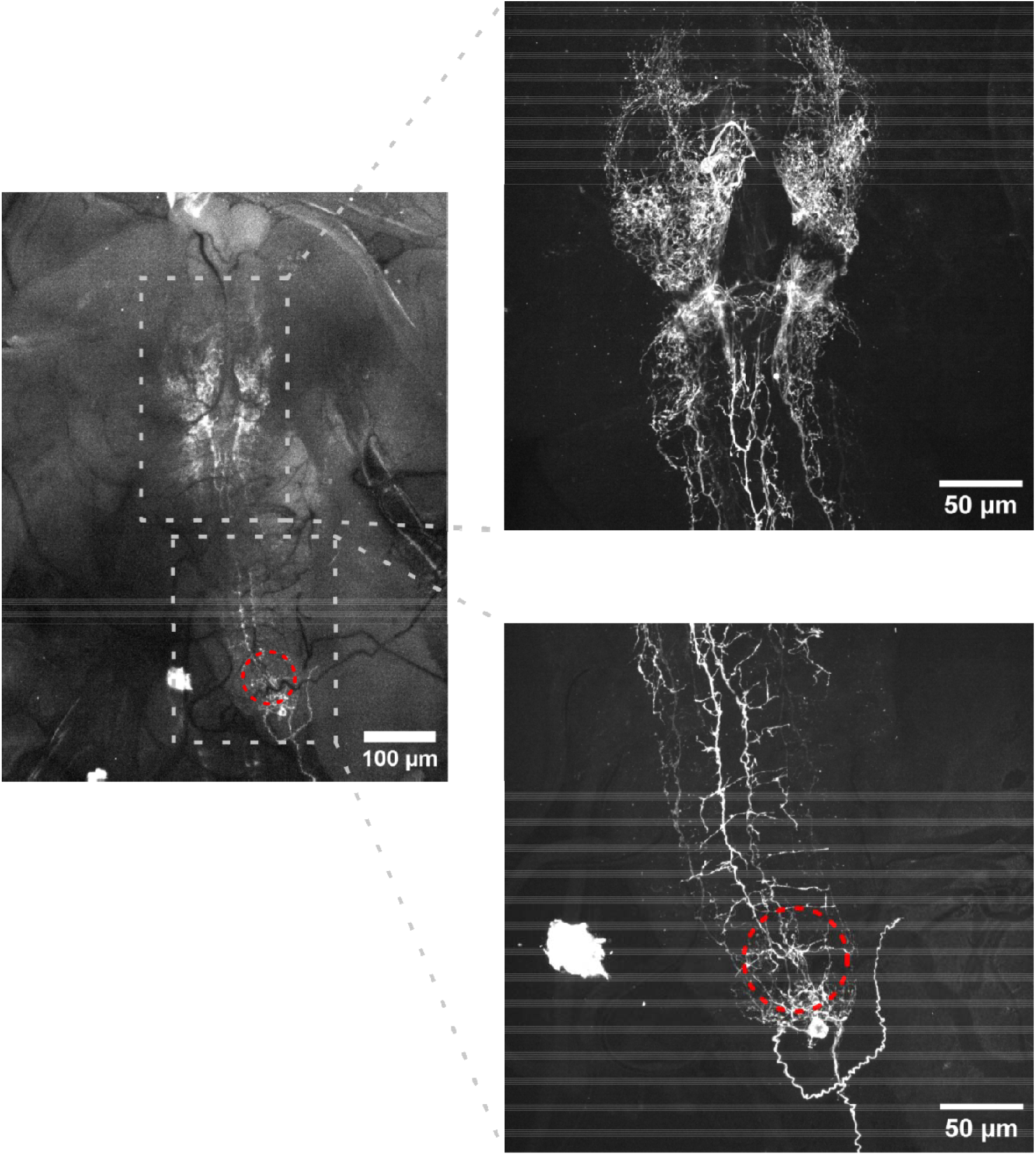
Restricted *tdc*^*2+*^ Neuron Anatomy to the Brain. Confocal staining of 20XUAS-CsChrimson-venus in attP18;tsh-LexA, pJFRC20-8XLexAop2-IVS-GAL80-WPRE attP40/ CyO, Tb-RFP; MKRS/TM6B with w[*]; P{w[±mC]=Tdc2-GAL4 at 10x magnification (left) and 40x magnification (right) of brain-residing *tdc*^*2+*^ neurons. Note the absence of soma within the VNC (left bottom) and the extensive descending projections from soma located in the SOG (left top). Red dashed-circle represent the focal point of restricted optogenetic stimulation.

## Notes

https://doi.org/10.17630/98cb0f9f-f064-4d08-addf-51d0025d1a7c

### Competing Interest Statement

The authors have declared no competing interest.

